# Intact synapse structure and function after combined knockout of PTPδ, PTPσ and LAR

**DOI:** 10.1101/2021.01.17.427005

**Authors:** Javier Emperador-Melero, Giovanni de Nola, Pascal S. Kaeser

**Affiliations:** Department of Neurobiology, Harvard Medical School, Boston, MA 02115

## Abstract

It has long been proposed that Leukocyte common Antigen-Related Receptor Protein Tyrosine Phosphatases (LAR-RPTPs) are cell-adhesion proteins for the control of synapse assembly. Their synaptic nanoscale localization, however, has not been established, and the fine structure of synapses after knockout of the three vertebrate genes for LAR-RPTPs (PTPδ, PTPσ and LAR) has not been tested. Here, we find that PTPδ is precisely apposed to postsynaptic scaffolds at excitatory and inhibitory synapses using superresolution microscopy. We generated triple-conditional knockout mice for PTPδ, PTPσ and LAR to test whether they are essential for synapse structure. While mild effects on synaptic vesicle clustering and active zone architecture were detected, synapse numbers and their overall structure were unaffected, membrane anchoring of the active zone persisted, and vesicle docking and release were normal. We conclude that LAR-RPTPs, despite their localization at synaptic appositions, are dispensable for the organization and function of presynaptic nerve terminals.

## Introduction

Presynaptic nerve terminals are packed with neurotransmitter-laden vesicles that fuse at the active zone, membrane-attached protein machinery that forms vesicular release sites. Work over the past two decades has established that the unique synaptic architecture with nanoscale apposition of these secretory hotspots with receptors on postsynaptic cells allows for robust signal transmission (Biederer et al., 2017; Südhof, 2012). The assembly mechanisms of these transcellular molecular machines, however, remain largely obscure (Emperador-Melero and Kaeser, 2020; Rizalar et al., 2021; Südhof, 2018).

Leukocyte common Antigen-Related Receptor Protein Tyrosine Phosphatases (LAR-RPTPs) are transmembrane proteins often regarded as master presynaptic organizers. Three LAR-RPTPs, PTPδ, PTPσ and LAR, are expressed in the brain, bind to the active zone organizer Liprin-α and to synaptic cell-adhesion proteins, and recruit presynaptic material in artificial synapse formation assays (Bomkamp et al., 2019; Han et al., 2018, 2020a; Johnson and Van Vactor, 2003; Kwon et al., 2010; Pulido et al., 1995; Serra-Pages et al., 1998; Takahashi et al., 2011; Yim et al., 2013). While these data suggest roles in presynaptic assembly (Fukai and Yoshida, 2020; Takahashi and Craig, 2013; Um and Ko, 2013), LAR-RPTP localization and function at neuronal synapses are less clear. In invertebrates, loss-of-function mutations in LAR-RPTPs resulted in defects in axon guidance, increased active zone and synapse areas, and impaired neurotransmitter secretion (Ackley et al., 2005; Clandinin et al., 2001; Desai et al., 1997; Kaufmann et al., 2002; Krueger et al., 1996). In mice, RNAi-mediated knockdown of LAR-RPTPs or deletion of PTPσ caused generalized loss of synapse markers and defective synaptic transmission (Dunah et al., 2005; Han et al., 2018, 2020a, 2020b), leading to the model that LAR-RPTPs control synapse formation. Furthermore, mild synaptic and behavioral defects were observed in single gene constitutive knockouts (Elchebly et al., 1999; Horn et al., 2012; Park et al., 2020; Uetani et al., 2000; Wallace et al., 1999). Contrasting the RNAi-based analyses, however, a recent study used conditional mouse gene targeting to ablate PTPδ, PTPσ and LAR, and found no overt defects in neurotransmitter release (Sclip and Südhof, 2020), thereby questioning the general role of LAR-RPTPs in synapse assembly.

The lack of knowledge of LAR-RPTP nanoscale localization and of a characterization of vertebrate synapse structure after ablation of all LAR-RPTPs obscures our understanding of their roles as synapse organizers. Here, we establish that PTPδ is apposed to postsynaptic scaffolds of inhibitory and excitatory synapses using stimulated emission depletion (STED) microscopy, supporting that these proteins could control synapse formation or regulate synapse function. However, analyses of active zone protein composition, synapse ultrastructure, and synaptic transmission in newly generated conditional PTPδ/PTPσ/LAR triple knockout mice reveal that these proteins are largely dispensable for synapse structure and function.

## Results

PTPδ, PTPσ and LAR are encoded by *Ptprd, Ptprs* and *Ptprf*, respectively. Conditional knockout mice for each gene were generated using homologous recombination (Fig. S1). Alleles for PTPδ (Farhy-Tselnicker et al., 2017; Sclip and Südhof, 2020) and PTPσ (Bunin et al., 2015; Sclip and Südhof, 2020) were identical to previously reported alleles, while the LAR allele was newly generated. The floxed alleles for each gene did not impair survival or RPTP protein expression (Fig. S1). We intercrossed these alleles to generate triple-conditional knockout mice. In cultured hippocampal neurons, Cre recombinase was delivered by lentiviruses and expressed from a Synapsin promoter (Liu et al., 2014), and resulted in removal of PTPδ, PTPσ and LAR, generating cTKO^RPTP^ neurons (Figs. 1A, 1B). Control^RPTP^ neurons were obtained using an inactive version of Cre.

**Figure 1.**
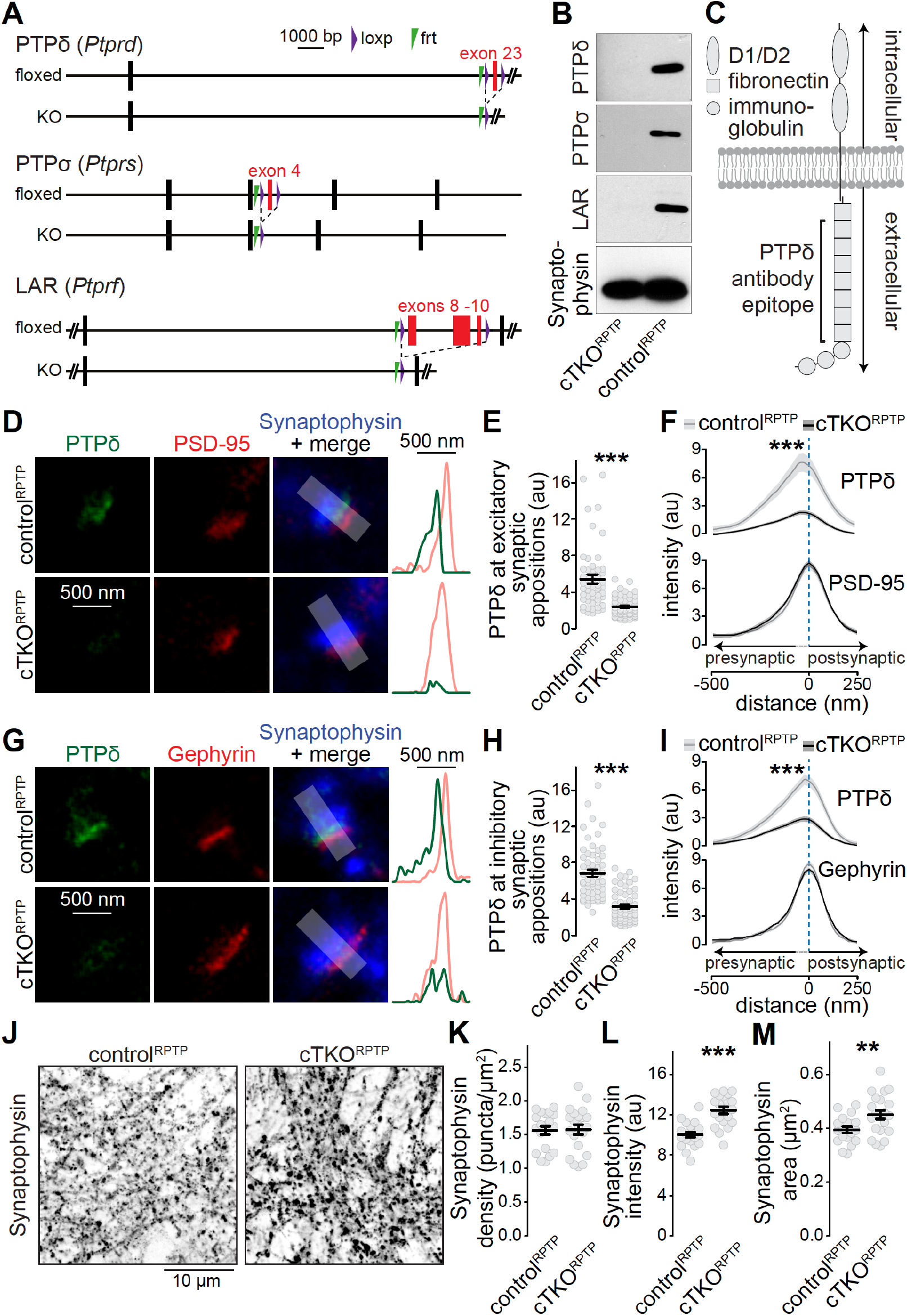
Triple conditional LAR-RPTP knockout to assess nanoscale localization of PTPδ and roles of LAR-RPTP in synapse formation. **(A)** Diagram for simultaneous conditional knockout of PTPδ, PTPσ and LAR by cre recombination. **(B)** Example western blot of cultured neurons from PTPδ, PTPσ and LAR triple floxed mice expressing Cre recombinase (to generate cTKO^RPTP^ neurons) or truncated Cre (to generate control^RPTP^ neurons). The bands detected in the cultured neurons correspond to the lower bands detected in brain homogenate shown in Fig. S1. **(C)** Diagram showing the general structure of LAR-RPTPs and the antibody recognition site for PTPδ (antibodies were generated using a peptide containing fibronectin domains 2, 3 and 8 (Shishikura et al., 2016)). **(D-F)** STED images, (D) quantification of the peak intensity of PTPδ (E) and average intensity profiles for PTPδ and PSD-95 (F) at single excitatory synapses. Side-view synapses were identified by the presence of bar-like PSD-95 signals at the edge of the vesicle cloud marked by Synaptophysin. Intensity profiles of shaded areas in the overlap images were used to determine the peak intensity of the protein of interest, and are shown on the right of the corresponding image. N (control^RPTP^) = 50 synapses/3 cultures, N (cTKO^RPTP^) = 54/3. **(G-I)** Same as D-F, but for inhibitory synapses identified by Gephyrin. N (control^RPTP^) = 58/3 cultures, N (cTKO^RPTP^) = 59/3. **(J-M)** Confocal images of cultured neurons stained with anti-Synaptophysin antibodies (J) and quantification of Synaptophysin puncta density (K), intensity (L) and size (M) detected using automatic two-dimensional segmentation. N (control^RPTP^) = 20 images/3 independent cultures; N (cTKO^RPTP^) = 21/3. The Synaptophysin confocal data are from the experiments shown in D-I. Data are plotted as mean ± SEM and were analyzed using two-way ANOVA tests (F, I, genotype *** for PTPδ), t-tests (E, L, M) or Mann-Whitney rank sum tests (H, K). ** p < 0.01, *** p < 0.001.

We first aimed at resolving the subsynaptic localization of LAR-RPTPs using STED microscopy. PTPδ antibody specificity was established using cTKO^RPTP^ neurons as controls, while antibodies suitable for superresolution analyses of PTPσ or LAR could not be identified. To determine PTPδ localization, we selected side-view synapses with bar-like postsynaptic receptor scaffolds (PSD-95 and Gephyrin for excitatory and inhibitory synapses, respectively) on one side of a Synaptophysin-labeled nerve terminal (Fig. S2, (Emperador-Melero et al., 2020; Held et al., 2020; Wong et al., 2018)). PTPδ, detected with antibodies against the extracellular fibronectin domains (Shishikura et al., 2016), was concentrated apposed to PSD-95 and Gephyrin, respectively, at distances of 24 ± 17 nm (PSD-95) and 28 ± 11 nm (Gephyrin) (Figs. 1D-1I). Only background signal typical for quantification of raw images (Emperador-Melero et al., 2020; Held et al., 2020; Wang et al., 2016; Wong et al., 2018) remained in cTKO^RPTP^ neurons in STED (Figs. 1D-1I) and confocal (Fig. S3) microscopy. This establishes that the extracellular portion of PTPδ localizes to the synaptic cleft. Given the presynaptic roles in invertebrate synapses and synapse formation assays (Ackley et al., 2005; Kaufmann et al., 2002), the interactions with the active zone protein Liprin-α (Pulido et al., 1995; Serra-Pages et al., 1998; Wong et al., 2018), and the asymmetry of the average STED side-view profile with a bias towards the presynapse (Figs. 1F, 1I), we conclude that most PTPδ is presynaptic and localized at the active zone, but postsynaptic components cannot be excluded. Furthermore, most synapses contain PTPδ, as 88% of excitatory and 92% of inhibitory synapses had PTPδ peak intensities higher than three standard deviations above the average of the cTKO^RPTP^ signal.

The subsynaptic PTPδ localization and its presence at most synapses is consistent with general roles of LAR-RPTPs in synapse organization. However, the synapse density, measured as Synaptophysin puncta, was unchanged in cTKO^RPTP^ neurons (Figs. 1L-1O), indicating that LAR-RPTPs are not necessary for synapse formation. Small increases in puncta intensity and area were detected (Figs. 1L-1O), consistent with enlargements observed in invertebrates (Ackley et al., 2005; Kaufmann et al., 2002). A recent independent study that ablated LAR-RPTPs early also found normal synapse densities (Sclip and Südhof, 2020), contradicting the generalized model that LAR-RPTPs are master synapse organizers (Dunah et al., 2005; Fukai and Yoshida, 2020; Han et al., 2018, 2020a, 2020b; Kwon et al., 2010; Takahashi and Craig, 2013; Um and Ko, 2013; Yim et al., 2013). It remains possible that LAR-RPTPs control assembly of a specific subset of synapses, which may explain why PTPδ ablation causes modest layer-specific impairments of synaptic strength (Park et al., 2020).

We next examined whether LAR-RPTPs have specific roles in presynaptic nanoscale structure. Electron microscopy of high-pressure frozen neurons (Figs. 2A-2E) revealed that synaptic vesicles were efficiently clustered at cTKO^RPTP^ synapses. A ~15% increase in the total synaptic vesicle number per synapse profile was detected, matching the modestly increased Synaptophysin signals (Fig. 1) and the enhanced presence of vesicular markers in C.elegans mutants (Ackley et al., 2005). Notably, no differences in vesicle docking (defined by vesicles for which the electron dense membrane merges with the electron density of the target membrane) were observed. Synapse width, quantified as the width of the synaptic cleft, was increased by ~30%, again matching invertebrate phenotypes (Kaufmann et al., 2002). These data establish that LAR-RPTP ablation does not strongly impair synapse ultrastructure. LAR-RPTPs may shape aspects of the synaptic cleft, consistent with their localization and transsynaptic interactions and possibly similar to other synaptic cell adhesion proteins, for example SynCAMs (Perez de Arce et al., 2015).

**Figure 2.**
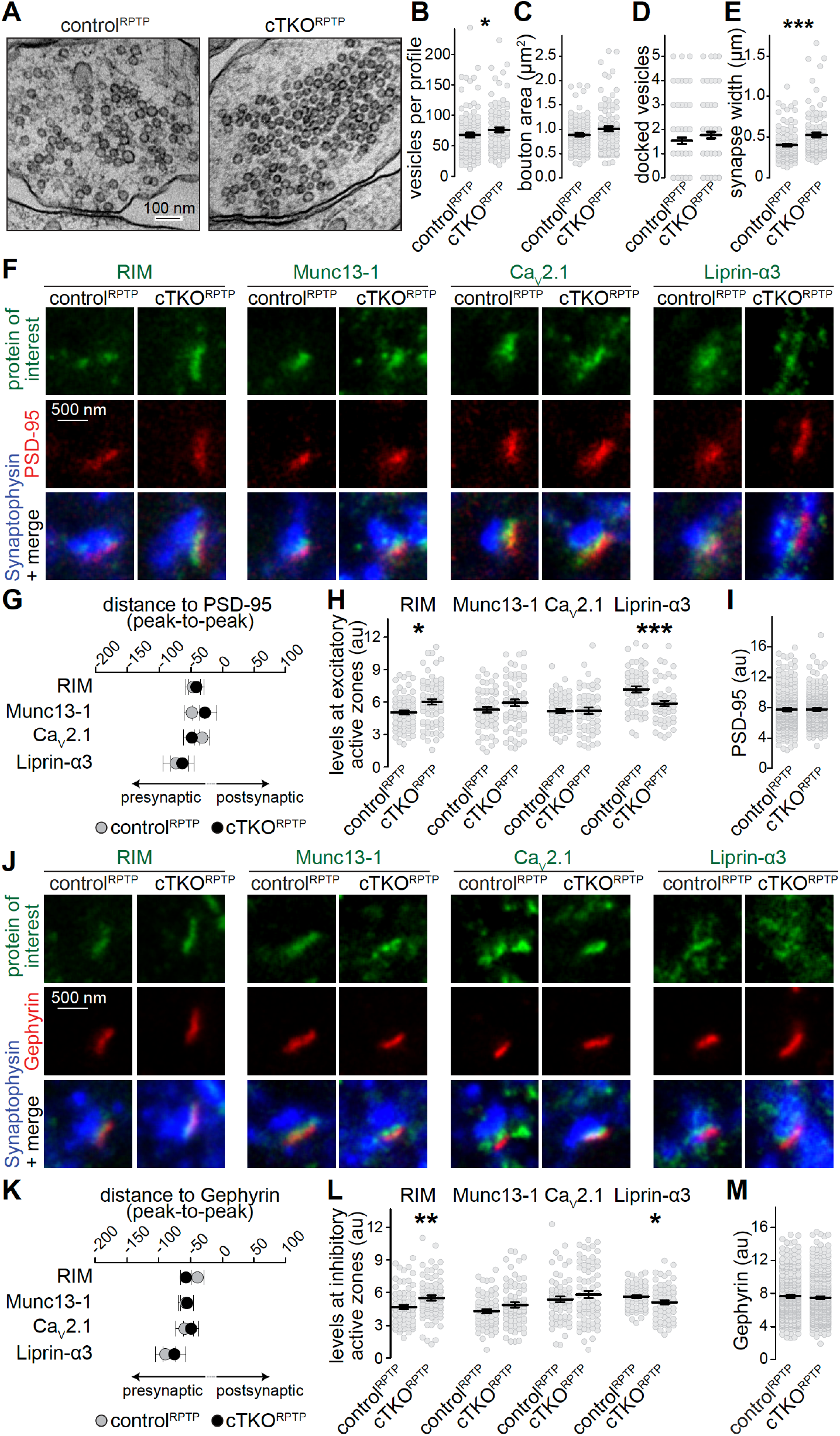
Synapse ultrastructure and active zone composition after LAR-RPTP knockout. **(A-E)** Electron micrographs (A) and quantification of the total number of vesicles per profile (B), bouton area (C), number of docked vesicles (D) and synapse width (E) assessed in single sections of high-pressure frozen neurons. N (control^RPTP^) = 106 synapses/2 independent cultures, N (cTKO^RPTP^) = 101/2. **(F-H)** STED example images of excitatory side-view synapses (F) and quantification of the distance to PSD-95 (G) and peak intensities (H) of RIM, Munc13-1, Ca_V_2.1 and Liprin-α3. RIM: N (control^RPTP^) = 68 synapses/3 independent cultures, N (cTKO^RPTP^) = 68/3; Munc13-1: N (control^RPTP^) = 57/3, N (cTKO^RPTP^) = 60/3; Ca_V_2.1: N (control^RPTP^) = 64/3, N (cTKO^RPTP^) = 58/3; Liprin-α3: N (control^RPTP^) = 56/3, N (cTKO^RPTP^) = 53/3. **(I)** Quantification of the peak intensity of PSD-95. N (control^RPTP^) = 295/3; N (cTKO^RPTP^)= 293/3. (**J-L**) Same as F-H, but for Gephyrin-containing inhibitory synapses. RIM: N (control^RPTP^) = 75/3 cultures, N (cTKO^RPTP^) = 79/3; Munc13-1: N (control^RPTP^) = 65/3, N (cTKO^RPTP^) = 72/3; Ca_V_2.1: N (control^RPTP^) = 64/3, N (cTKO^RPTP^) = 71/3; Liprin-α3: N (control^RPTP^) = 65/3, N (cTKO^RPTP^) = 61/3. **(M)** Quantification of the peak intensity of Gephyrin. N (control^RPTP^) = 327/3; N (cTKO^RPTP^) = 342/3. Data are plotted as mean ± SEM and were analyzed using Mann-Whitney rank sum tests. * p < 0.05, ** p < 0.01, *** p < 0.001.

We assessed whether active zone proteins, which are present at normal levels in Western blots after LAR-RPTP ablation (Sclip and Südhof, 2020), are anchored at the presynaptic membrane by LAR-RPTPs. STED microscopy was used to measure localization and peak levels of active zone proteins at excitatory (Figs. 2F-2I) and inhibitory (Figs. 2J-2M) synapses. RIM, Munc13-1, Ca_V_2.1 and Liprin-α3 were localized within ~30-~60 nm of the postsynaptic scaffolds in control^RPTP^ and cTKO^RPTP^ synapses, as expected for these proteins (Held et al., 2020; Wong et al., 2018). Overall, there were no strong changes in their levels, but small increases in RIM and small decreases in Liprin-α3 were detected in both types of cTKO^RPTP^ synapses either by STED (Figs. 2F-2M) or confocal (Fig. S4) microscopy. While binding between Liprin-α and LAR-RPTPs (Pulido et al., 1995; Serra-Pages et al., 1998) may explain Liprin-α3 reductions, these data establish that other pathways are sufficient to recruit most Liprin-α3 to active zones. The higher levels of RIM may be compensatory to reductions in Liprin-α3, and could be related to the liquid-liquid phase separation properties of both proteins (Emperador-Melero et al., 2020; McDonald et al., 2020; Wu et al., 2019). Overall, we conclude that the active zone remains assembled and anchored to the target membrane in the absence of LAR-RPTPs.

A previous study found that LAR-RPTP ablation induced no strong defects in glutamate release, but regulated NMDARs through a transsynaptic mechanism (Sclip and Südhof, 2020). These findings are consistent with the near-normal synaptic ultrastructure and active zone assembly (Fig. 2). We complemented this recent study by whole-cell recordings of inhibitory postsynaptic currents (IPSCs, Fig. 3), which were not previously assessed after LAR-RPTP ablation. Release evoked by single action potentials was similar between control^RPTP^ and cTKO^RPTP^ neurons and IPSC kinetics were unaffected. The IPSC ratio of two consecutive stimuli (paired pulse ratio), which is inversely proportional to vesicular release probability (Zucker and Regehr, 2002), was also unaffected. We conclude that synaptic vesicle exocytosis, here monitored via IPSCs, is not impaired by LAR-RPTP knockout.

**Figure 3.**
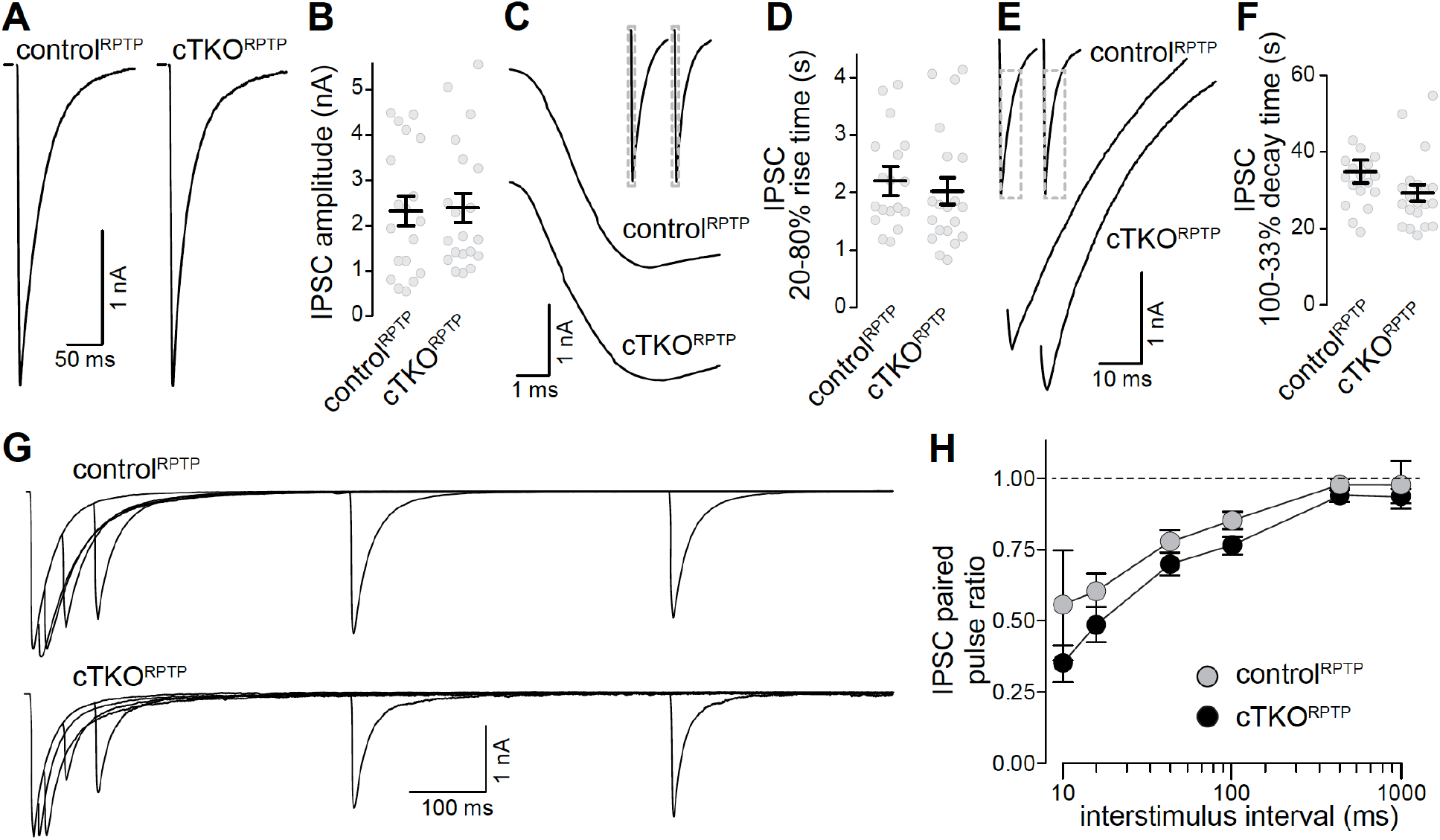
Synaptic transmission in LAR-RPTP triple knockout neurons. **(A, B)** Example traces (A) and average amplitudes (B) of single action potential evoked IPSCs. N (control^RPTP^) = 19 cells/3 independent cultures, N (cTKO^RPTP^) = 20/3. **(C, D)** Example zoomed-in trace of the IPSC rise (C) and quantification of 20-80% rise times (D) of evoked IPSCs, N as in A, B. **(E, F)** Example zoomed-in trace of the IPSC decay (E) and quantification of 100-33% decay times (F) of evoked IPSCs. N as in A, B. **(G, H)** Example traces (G) and average IPSC paired pulse ratios (H) at various interstimulus intervals. N (control^RPTP^) = 18/3, N (cTKO^RPTP^) = 19/3. Data are plotted as mean ± SEM and were analyzed using Mann-Whitney rank sum tests (B, D, F) or a two-Way ANOVA (H), no significant differences were detected.

## Discussion

Overall, we demonstrate that ablation of LAR-RPTPs from vertebrate synapses does not alter synapse density, vesicle docking, membrane anchoring of active zones, and synaptic vesicle release. This aligns with a parallel study that reported no loss of synaptic puncta and efficient release at excitatory synapses in cultured hippocampal neurons and in acute hippocampal brain slices (Sclip and Südhof, 2020) upon LAR-RPTP knockout, but contrasts RNAi-based studies that led to models in which these RPTPs are major synapse organizers (Dunah et al., 2005; Fukai and Yoshida, 2020; Han et al., 2018, 2020a, 2020b; Kwon et al., 2010; Takahashi and Craig, 2013; Um and Ko, 2013; Yim et al., 2013). LAR-RPTPs belong to the superfamily of RPTPs (Johnson and Van Vactor, 2003), and it is possible that different RPTPs compensate for their loss. We note, however, that the time course of deletion in our knockout experiments is similar to the time course that is used in most RNAi-knockdown studies, and is hence unlikely to explain the differences. Other contributing factors could be different experimental preparations and off-target effects of knockdowns, which may generate artifacts in synapse formation experiments (Südhof, 2018). Altogether, we conclude that, while biochemical and synapse formation assays support synaptogenic activities for these proteins, synapses persist upon LAR-RPTP ablation, and their structure and function do not necessitate these proteins.

Our study establishes specific localization of PTPδ extracellular domains to the synaptic cleft. Hence, PTPδ is correctly positioned to locally execute synaptic functions, for example for shaping cleft geometry, to modulate presynaptic plasticity, or to control postsynaptic receptors (Biederer et al., 2017; Sclip and Südhof, 2020; Uetani et al., 2000). Such functions would not be at odds with the relatively mild structural and functional effects after LAR-RPTP ablation, nor with upstream functions in neurite outgrowth and axon targeting (Ackley et al., 2005; Clandinin et al., 2001; Desai et al., 1997; Krueger et al., 1996; Prakash et al., 2009; Shishikura et al., 2016). Mechanisms of active zone anchoring to the target membrane, however, remain unresolved. Deletion of the major candidates, Ca_V_2 channels (Held et al., 2020), Neurexins (Chen et al., 2017), and now LAR-PTPs (Figs. 1, 2) produce no major structural defects, indicating that active zones are most likely anchored to the plasma membrane through multiple parallel pathways (Emperador-Melero and Kaeser, 2020).

## Acknowledgements

We thank J. Wang, M. Han and C. Qiao for technical support, Dr. H. Nyitrai for help and advice, Dr. F. Nakamura for PTPδ antibodies, and Drs. M. Verhage and J. Broeke for the SynapseEM MATLAB macro. This work was supported by grants from the NIH (R01NS083898 and R01MH113349 to PSK), the Lefler Foundation (to PSK), the Armenise Harvard Foundation (to PSK), and a fellowship from the Alice and Joseph E. Brooks postdoctoral fund (to JEM). For mutant mice, we thank the Welcome Trust Sanger Institute Mouse Genetics Project (Sanger MGP), its funders and INFRAFRONTIER/EMMA (www.infrafrontier.eu, *Ptprd*, funding information may be found at www.sanger.ac.uk/mouseportal and associated primary phenotypic information at www.mousephenotype.org), the Canadian Mouse Mutant Repository at the Hospital for Sick Children (*Ptprs*), and the Helmholtz Zentrum München (*Ptprf*). We thank the Dana-Farber/Harvard Cancer Center for the use of the Transgenic Mouse Core (in part supported by an NCI Cancer Center Support Grant # P30CA006516), which performed cryo-recovery, embryonic stem cell expansion, and blastocyst injections. We acknowledge the Neurobiology Imaging Facility (supported by a P30 Core Center Grant NS072030), and the Electron Microscopy Facility at Harvard Medical School.

## Author contributions

Conceptualization, J.E-M. and P.S.K.; Methodology, J.E-M. and G.dN.; Investigation, J.E-M. and G.dN.; Formal Analysis, J.E-M., G.dN. and PSK; Writing-Original Draft, J.E-M and P.S.K.; Supervision, P.S.K.; Funding Acquisition, P.S.K.

## Materials and methods

### Mouse lines

PTPδ (*Ptprd*) mice were acquired as frozen embryos from the Welcome Trust Sanger Institute (Ptprd^tm2a(KOMP)Wtsi^; clone EPD0581_9_D04, MGI:4458607, RRID:IMSR_EM:11805) and the same mutant allele was described in previous studies (Farhy-Tselnicker et al., 2017; Sclip and Südhof, 2020). PTPσ (*Ptprs*) mice were obtained as frozen sperm from the Canadian Mouse Mutant Repository at the Hospital for Sick Children (C57BL/6N-Ptprs^tm1a(KOMP)Mbp^/Tcp; clone DEPD00535_1_D11; MGI:4840831, RRID:IMSR_CMMR:ABCA), and were also used previously (Bunin et al., 2015; Sclip and Südhof, 2020). Embryonic stem cells containing the LAR (*Ptprf*) mutant allele were obtained from the Helmholtz Zentrum München (Ptprf^tm1a(EUCOMM)Wtsi^; clone EPD0697_1_D03; MGI:4887720). Mutant alleles were originally generated using homologous recombination by the international knockout consortium (Bradley et al., 2012; Skarnes et al., 2011). Frozen embryos (PTPδ), frozen sperm (PTPσ) or embryonic stem cells (LAR) were used to establish the respective mouse lines through the Transgenic Mouse Core (DF/HCC) at Harvard Medical School. For generation of the LAR mutant mice, the embryonic stem cells were expanded, the genotype was confirmed by PCR and sequencing, and injection into C57BL/6 blastocysts was used to generate chimeric founders. After germline transmission, the mice were crossed to Flp-expressing mice (Dymecki, 1996) to remove the LacZ and Neomycin cassettes to generate the conditional allele. The same crossing was performed with the cryo-recovered PTPδ and PTPσ mice. This strategy generated conditional “floxed” alleles for each gene, in which exon 23 for *Ptprd*, exon 4 for *Ptprs*, and exons 8, 9 and 10 for *Ptprf* were flanked by loxP sites. Survival of each individual floxed allele was analyzed in offsprings of heterozygote matings through comparison of obtained genotypes of offsprings on or after P14 to expected genotypes for Mendelian inheritance. The three floxed lines were intercrossed and maintained as triple-homozygote mice. The conditional PTPδ, PTPσ and LAR alleles were genotyped using the oligonucleotide primers CAGAGGTGGCTCATGTGC and GCCCAACCCTCAATTGTCAGAC (PTPδ, 465 and 287 bp bands for the floxed and wild-type alleles, respectively), GAGTCCTCAAACCAGGCCCTG and GGTGAGACCAGGGTGGGTTC (PTPσ, 522 and 345 bp bands for the floxed and wild-type alleles, respectively), and CAGAGGTGGCTCATGTGC and GCCCAACCCTCAATTGTCAGAC (LAR, 498 and 289 bp bands for the floxed and wild-type alleles, respectively). All animal experiments were approved by the Harvard University Animal Care and Use Committee.

### Neuronal cultures and production of lentiviruses

Primary hippocampal cultures were prepared as described (Emperador-Melero et al., 2020; Held et al., 2020; Wong et al., 2018). Briefly, hippocampi of newborn (postnatal days P0 or P1) pups were digested in papain, and neurons were plated onto glass coverslips in Plating Medium composed of Mimimum Essential Medium (MEM) supplemented with 0.5% glucose, 0.02% NaHCO_3_, 0.1 mg/ml transferrin, 10% Fetal Select bovine serum, 2mM L-glutamine, and 25 mg/ml insulin. After 24 h, Plating Medium was exchanged with Growth Medium composed of MEM with 0.5% glucose, 0.02% NaHCO_3_, 0.1 mg/ml transferrin, 5% Fetal Select bovine serum (Atlas Biologicals), 2% B-27 supplement, and 0.5 mM L-glutamine. At DIV2-3, Cytosine b-D-arabinofuranoside (AraC) was added to a final concentration of 1 to 2 mM. Cultures were kept in a 37 °C incubator for a total of 14 to 16 d before analyses proceeded. Lentiviruses were produced in HEK293T cells maintained in DMEM supplemented with 10% bovine serum and 1% penicillin/streptomycin. HEK293T cells were transfected using calcium phosphate precipitation with a combination of three lentiviral packaging plasmids (REV, RRE and VSV-G) and a separate plasmid encoding either Cre recombinase or inactive Cre, at a molar ratio of 1:1:1:1. 24 h after transfection, the medium was changed to neuronal growth medium and 18 - 30 h later the supernatant was used for viral transduction. Neuronal cultures were infected 6 d after plating with lentiviruses expressing GFP-Cre or an inactive variant of GFP-Cre expressed under the human Synapsin promotor (Liu et al., 2014), and infection rates were assessed via nuclear GFP. Only cultures in which no non-infected neurons could be detected were used for analyses.

### Western blotting

Cell lysates were collected from DIV14-15 neuronal cultures in a 1x sodium dodecyl sulfate (SDS) solution in PBS. For tissue collection, brains of postnatal day P21 to P28 mice were homogenized using a glass-Teflon homogenizer in 5 ml of ice-cold homogenizing solution (150 mM NaCl, 25 mM HEPES, 4 mM EDTA and 1% Triton X-100, pH 7.5), following addition of SDS (to a final concentration of 1x). All samples were denatured at 100 °C for 10 min, run on SDS-PAGE gels, and then transferred to nitrocellulose membranes for 6.5 h at 4 °C in buffer containing (per l) 200 ml methanol, 14 g glycine and 6 g Tris. Next, membranes were blocked for 1 h at room temperature in saline buffer with 10% non-fat milk powder and 5% normal goat serum. Membranes were incubated in primary antibodies overnight at 4 °C in PBS with 5% milk and 2.5% goat serum, followed by 1 h incubation with horseradish peroxidase (HRP)-conjugated secondaries at room temperature. Three 5 min washes were performed between every step. Protein bands were visualized using chemiluminescence and exposure to film. The primary antibodies were: goat anti-PTPσ (A114, 1:200, RRID: AB_2607944), rat anti-PTPδ (A229, 1:500, gift of Dr. F. Nakamura (Shishikura et al., 2016)), mouse anti-LAR (A156, 1:500, clone E9B9S from Cell signaling), and mouse anti-Synaptophysin (A100, 1: 5000, RRID: AB_887824). For PTPσ, normal goat serum was substituted by rabbit serum. The secondary antibodies were HRP-conjugated goat anti-mouse IgG (S44, 1:10000, RRID:AB_2334540), HRP-conjugated goat anti-rabbit IgG (S45, 1:10000, RRID:AB_2334589), HRP-conjugated goat anti-rat IgG (S46, 1:10000, RRID:AB_10680316), and HRP-conjugated anti-goat antibodies (S60, 1:10000).

### Immunofluorescence staining of neurons

Neurons grown on #1.5 glass coverslips were fixed at DIV15 in 4 % paraformaldehyde (PFA) for 10 min (except for staining with anti-Ca_v_2.1 antibodies, for which 2% PFA was used), followed by blocking and permeabilization in PBS containing 3% BSA/0.1% Triton X-100/PBS for 1 h at room temperature. Incubation with primary and secondary antibodies was performed overnight at 4 °C and for 1 h at room temperature, respectively. Samples were post-fixed in 4% PFA for 10 min and mounted onto glass slides using ProLong diamond mounting medium. Antibodies were diluted in blocking solution. Three 5 min washes with PBS were performed between steps. Primary antibodies used were: rabbit anti-Liprin-α3 (A232, 1:250; home-made (Emperador-Melero et al., 2020)), rabbit anti-RIM (A58, 1:500, RRID: AB_887774), mouse anti-PSD-95 (A149, 1:500; RRID: AB_10698024), mouse anti-Gephyrin (A8, 1:500; RRID:AB_2232546), guinea pig anti-Synaptophysin (A106, 1:500; RRID: AB_1210382), rabbit anti-Munc13-1 (A72, 1:500; RRID: AB_887733), rat anti-PTPδ (A229; 1:500; gift of Dr. F. Nakamura (Shishikura et al., 2016)), and rabbit anti-Ca_V_2.1 (A46, 1:500; RRID: AB_2619841). Secondary antibodies used: goat anti-rabbit Alexa Fluor 488 (S5; 1:500, RRID:AB_2576217), goat anti-mouse IgG1 Alexa Fluor 555 (S19, 1:500, RRID: AB_2535769), goat anti-mouse IgG2a Alexa Fluor 633 (S30, 1:500, RRID: AB_1500826), goat anti-guinea pig IgG Alexa Fluor 405 (S51, 1:500, RRID: RRID:AB_2827755).

### STED and confocal imaging

All images were acquired as described (Emperador-Melero et al., 2020; Held et al., 2020; Wong et al., 2018) using a Leica SP8 Confocal/STED 3X microscope equipped with an oil-immersion 100X 1.44-N.A objective, white lasers, gated detectors, and 592 nm and 660 and 770 nm depletion lasers. For every region of interest (ROI), quadruple color sequential confocal scans for Synaptophysin, PSD-95, Gephyrin and a protein of interest (RIM, Munc13-1, PTPδ, Liprin-α or Ca_V_2.1) were followed by triple-color sequential STED scans for PSD-95, Gephyrin and the protein of interest. Synaptophysin was only imaged in confocal mode because of depletion laser limitations. Identical settings were applied to all samples within an experiment. For analyses of synapse density, Synaptophysin signals were used to generate ROIs using automatic detection with a size filter of 0.4 - 2 μm^2^ (code available at https://github.com/kaeserlab/3DSIM_Analysis_CL and https://github.com/hmslcl/3D_SIM_analysis_HMS_Kaeser-lab_CL) and as described before (Emperador-Melero et al., 2020; Held et al., 2020; Liu et al., 2018). To measure synaptic levels of PTPδ, RIM, Munc13-1, Liprin-α3 and Ca_V_2.1 in confocal mode, a mask was generated in ImageJ using an automatic threshold in the Synaptophysin or the PSD-95 channel, and the levels were measured within that mask. For STED quantification, side-view synapses were selected while blind to the protein of interest. They were defined as synapses that contained a vesicle cluster (imaged in confocal mode) with a single bar-like Gephyrin or PSD-95 structure (imaged by STED) along the edge of the vesicle cluster. A 1 μm-long, 250-nm-wide profile was selected perpendicular to the postsynaptic density marker and across its center. The peak levels of the protein of interest were then measured as the maximum intensity of the line profile within 100 nm of the postsynaptic density marker peaks (estimated area based on Wong et al., 2018) after applying a 5-pixel rolled average. For side-view plots, line scans from individual side-view synapses were aligned to the peak of PSD-95 or Gephyrin after the 5-pixel rolling average was applied, and averaged across images. Only for representative images, a smooth filter was added, brightness and contrast were linearly adjusted, and images were interpolated to match publication standards. These adjustments were made identically for images within an experiment. All quantitative analyses were performed on original images without any processing, and all data were acquired and analyzed by an experimenter blind to genotype. For PTPδ STED analyses, synapses were considered PTPδ-positive if the peak intensity was higher than three standard deviations above the average of the cTKO^RPTP^ signal, assessed separately in each individual culture.

### High-pressure freezing and electron microscopy

Electron microscopy was performed as previously described (Held et al., 2020; Wang et al., 2016). Briefly, DIV15 neurons grown on 6 mm sapphire cover slips were frozen with a Leica EM ICE high pressure freezer in extracellular solution containing 140 mM NaCl, 5 mM KCl, 2 mM CaCl_2_, 2 mM MgCl_2_, 10 mM glucose, 10 mM Hepes, 20 μM CNQX, 50 μM AP5 and 50 μM picrotoxin (pH 7.4, ∼310 mOsm). Freeze-substitution was done in acetone containing 1% osmium tetroxide, 1% glutaraldehyde, and 1% H_2_O as follows: −90 °C for 5 h, 5 °C per h to −20 °C, −20 °C for 12 h, and 10 °C per hour to 20 °C. Samples were then infiltrated in epoxy resin, and baked at 60 °C for 48 h followed by 80 °C overnight. Next, sapphire coverslips were removed from the resin block by heat shock, and samples were sectioned at 50 nm with a Leica EM UC7 ultramicrotome. Sections were mounted on a nickel slot grid with a carbon coated formvar support film, and counterstained by incubation with 2% lead acetate solution for 10 s, followed by rinsing with distilled water. Samples were imaged with a JEOL 1200EX transmission electron microscope equipped with an AMT 2k CCD camera. Images were analyzed using SynapseEM, a MATLAB macro provided by Drs. M. Verhage and J. Broeke. Bouton area was measured by outlining the perimeter of each synapse profile. Docked vesicles were defined as vesicles touching the presynaptic plasma membrane opposed to the PSD, with the electron density of the vesicular membrane merging with that of the target membrane. Synapse width was measured as the area between synaptically apposed cells in which an evenly spaced cleft was present and associated with pre- and postsynaptic densities. All data were acquired and analyzed by an experimenter blind to the genotype.

### Electrophysiology

Electrophysiological recordings were performed as described before (Emperador-Melero et al., 2020; Held et al., 2020; Wang et al., 2016). Neurons were recorded at DIV15-16 in whole-cell patch-clamp configuration at room temperature in extracellular solution containing (in mM) 140 NaCl, 5 KCl, 1.5 CaCl_2_, 2 MgCl_2_, 10 HEPES (pH 7.4) and 10 Glucose, supplemented with 20 μM CNQX and 50 μM D-AP5 to block AMPA and NMDA receptors, respectively. Glass pipettes were pulled at 2.5 – 4 MΩ and filled with intracellular solutions containing (in mM) 40 CsCl, 90 K-Gluconate, 1.8 NaCl, 1.7 MgCl_2_, 3.5 KCl, 0.05 EGTA, 10 HEPES, 2 MgATP, 0.4 Na_2_-GTP, 10 phosphocreatine, CsOH and 4 mM QX314-Cl (pH 7.4). Neurons were clamped at −70 mV, series resistance was compensated to 4 – 5 MΩ, and recordings in which the uncompensated series resistance was >15 MΩ at any time during the experiment were discarded. Electrical stimulation was applied using a custom bipolar electrode made from Nichrome wire. A Multiclamp 700B amplifier and a Digidata 1550 digitizer were used for data acquisition, sampling at 10 kHz and filtering at 2 kHz. Data were analyzed using pClamp. The experimenter was blind during data acquisition and analyses.

### Statistics

Summary data are shown as mean ± SEM. Unless noted otherwise, significance was assessed using t-tests or Mann-Whitney U tests depending on whether assumptions of normality and homogeneity of variances were met (assessed using Shapiro or Levene’s tests, respectively). Two-way ANOVA tests on a 200 nm wide window centered around the PSD-95 peak were used for line profile analyses of STED data, and Chi-square tests were used to assess mouse survival ratios. All data were analyzed by an experimenter blind to the genotype. For each dataset, the specific tests used are stated in each figure legend.

---

**Figure S1.**
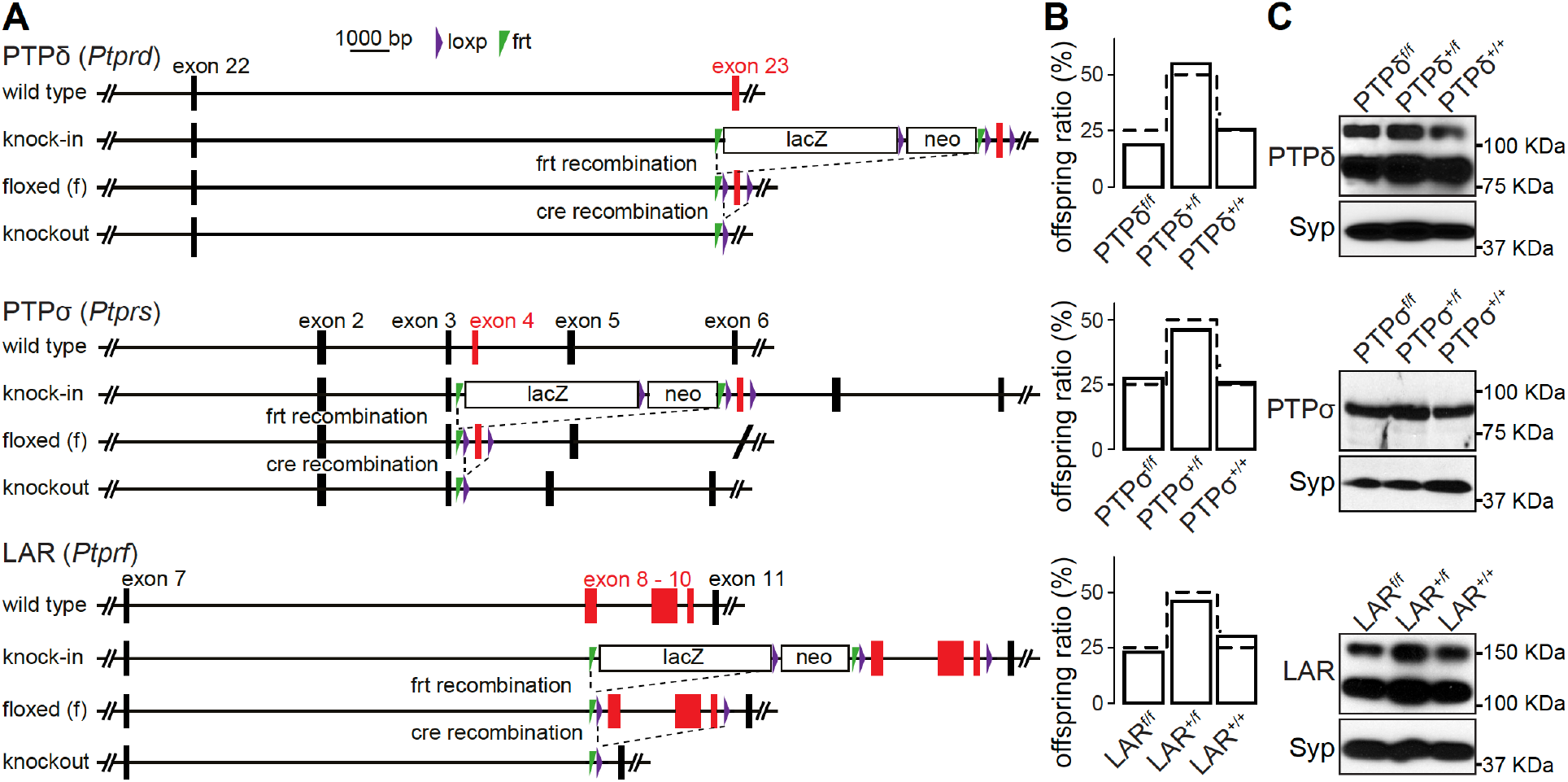
Generation of LAR-RPTP conditional knockout mice. **(A)** Gene targeting strategies for LAR-RPTP knockout mice. PTPδ and PTPσ alleles contain loxP sites flanking exons 23 and 4, respectively. They were imported for cryo-recovery at the “knock-in” stage from the Welcome Trust Sanger Institute (Ptprd^tm2a(KOMP)Wtsi^), and the Canadian Mouse Mutant Repository at the Hospital for Sick Children (C57BL/6N-Ptprs^tm1a(KOMP)Mbp^/Tcp), and are identical to the alleles described before (Bunin et al., 2015; Farhy-Tselnicker et al., 2017; Sclip and Südhof, 2020). The LAR allele contains loxp sites flanking exons 8, 9 and 10 and was obtained at the embryonic stem cell stage from the Helmholtz Zentrum München (Ptprf^tm1a(EUCOMM)Wtsi^). All three lines were crossed to flp-transgenic mice (Dymecki, 1996) to generate “floxed” conditional knockout alleles. **(B)** Survival analyses were performed on the offspring of heterozygote matings for each individual allele. Offspring ratios were assessed at >P14. Bars show the percentage of offspring for each genotype, and dotted lines represent the expected Mendelian ratio. N (PTPδ) = 28 mice/4 litters; N (PTPσ) = 31/4; N (LAR) = 22/3. Chi-square tests were used to compare expected Mendelian ratios with obtained offspring ratios, and no statistical difference was detected. **(C)** Western blots of whole brain homogenates of wild type, heterozygous and homozygous littermate mice for the PTPδ, PTPσ and LAR floxed alleles. For PTPδ, ~ 80 kDa and ~120 kDa bands were detected, matching the cleaved extracellular domains of the two main isoforms expressed in the brain containing 3 immunoglobulin and either 4 or 8 fibronectin domains (Shishikura et al., 2016). For PTPσ, a single band at ~ 90 KDa was detected, matching the size of the catalytically cleaved extracellular domain of the short isoform containing 4 fibronectin domains and 3 immunoglobulin domains (Aicher et al., 1997). For LAR, ~110 kDA and ~150 KDa bands were detected as previously described in the hippocampus (Yang et al., 2003). The bigger band matches the cleaved extracellular domains of the longest isoform containing 3 immunoglobulin and 8 fibronectin domains (Aicher et al., 1997), while the smaller band likely corresponds to a shorter isoform. In cultured hippocampal neurons, only the higher intensity bands at the lower molecular weight were detected, and these bands were effectively removed after cre-recombination of the floxed alleles (Fig. 1B).

**Figure S2.**
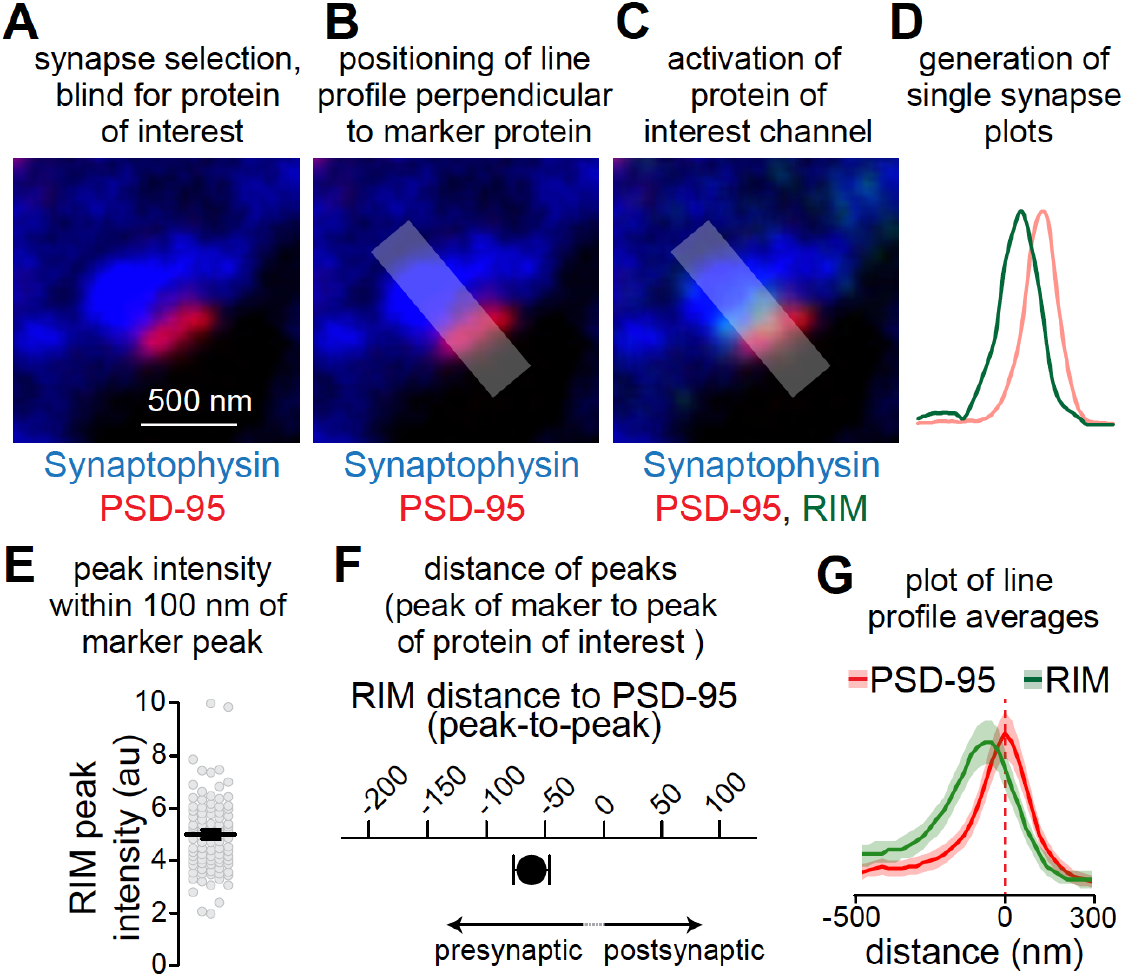
STED analysis workflow. (**A-D**) Workflow for STED analyses, showing an example of a side-view synapse immunostained for PSD-95 and RIM (both in STED mode) and Synaptophysin (in confocal mode). Side-view synapses are included when a PSD-95 bar is present at the edge of a Synaptophysin-labeled vesicle cloud (A). The synapse selection process is conducted by an experimenter blind for the protein of interest (RIM in the example shown). Next, a line profile is generated perpendicular to PSD-95 (B). The area for generating the line profile is shaded. The protein of interest channel is then activated (C) and the line profiles are generated (D). (**E-G**) Outline of the quantitative analyses across synapses. The peak intensity of the protein of interest within 100 nm of PSD-95 (Wong et al., 2018) is used to estimate protein levels in the synaptic cleft or at the active zone (E). The distance between RIM and PSD-95 peaks is used to estimate protein localization (F). The average of the line profiles of all synapses within an experiment is used to illustrate protein distribution (G). Data to illustrate STED workflow are from wild type hippocampal cultures, N = 42 synapses/2 independent cultures. Data are plotted as mean ± SEM.

**Figure S3.**
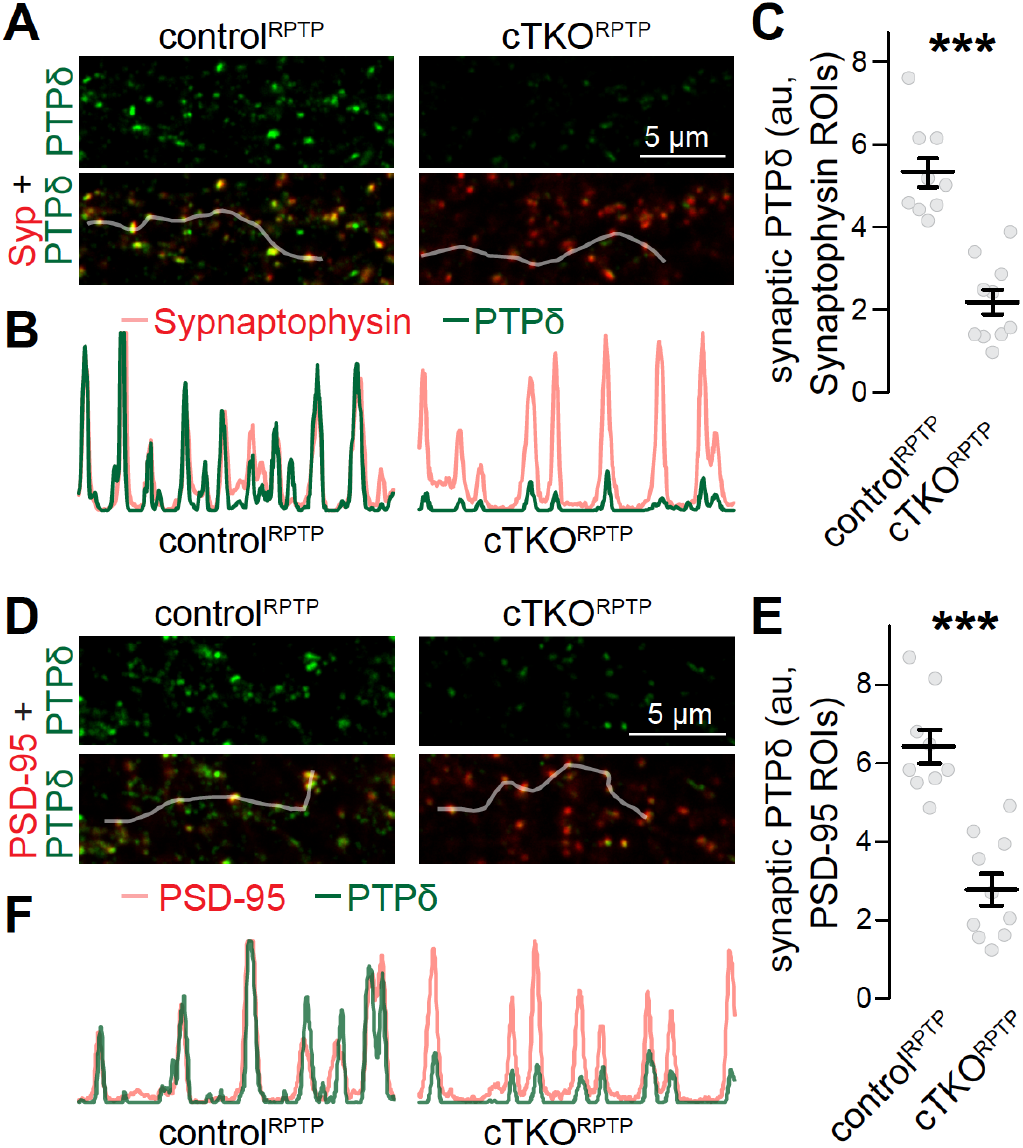
Confocal analyses of PTPδ. **(A-C)** Sample confocal images (A), sample intensity profiles (B) and quantification (C) of PTPδ at synapses identified by Synaptophysin (Syp) staining. Intensity profiles of PTPδ and Synaptophysin (C) along the shaded lines highlighted in A show good correlation between the signals. In C, PTPδ fluorescence intensities were quantified in Synaptophysin regions of interest (ROIs). The confocal images analyzed here were acquired in the same imaging session and for the same image frames as the STED analyses shown in Fig. 1. Confocal images were always acquired prior to STED acquisition, N (control^RPTP^) = 9 images/3 independent cultures; N (cTKO^RPTP^) = 10/3. **(D-F)** Same as A-C, but for PSD-95 ROIs. To avoid potential confounds of mildly increased Synaptophysin areas (Fig. 1M), we repeated the confocal analyses generating PSD-95 instead of Synaptophysin ROIs. In diffraction-limited microscopy, the resolution is insufficient to distinguish pre- and postsynaptic markers, and either marker can be used to generate synapse ROIs. N as in C. Data are plotted as mean ± SEM and analyzed using Mann-Whitney rank sum tests, *** p < 0.001.

**Figure S4.**
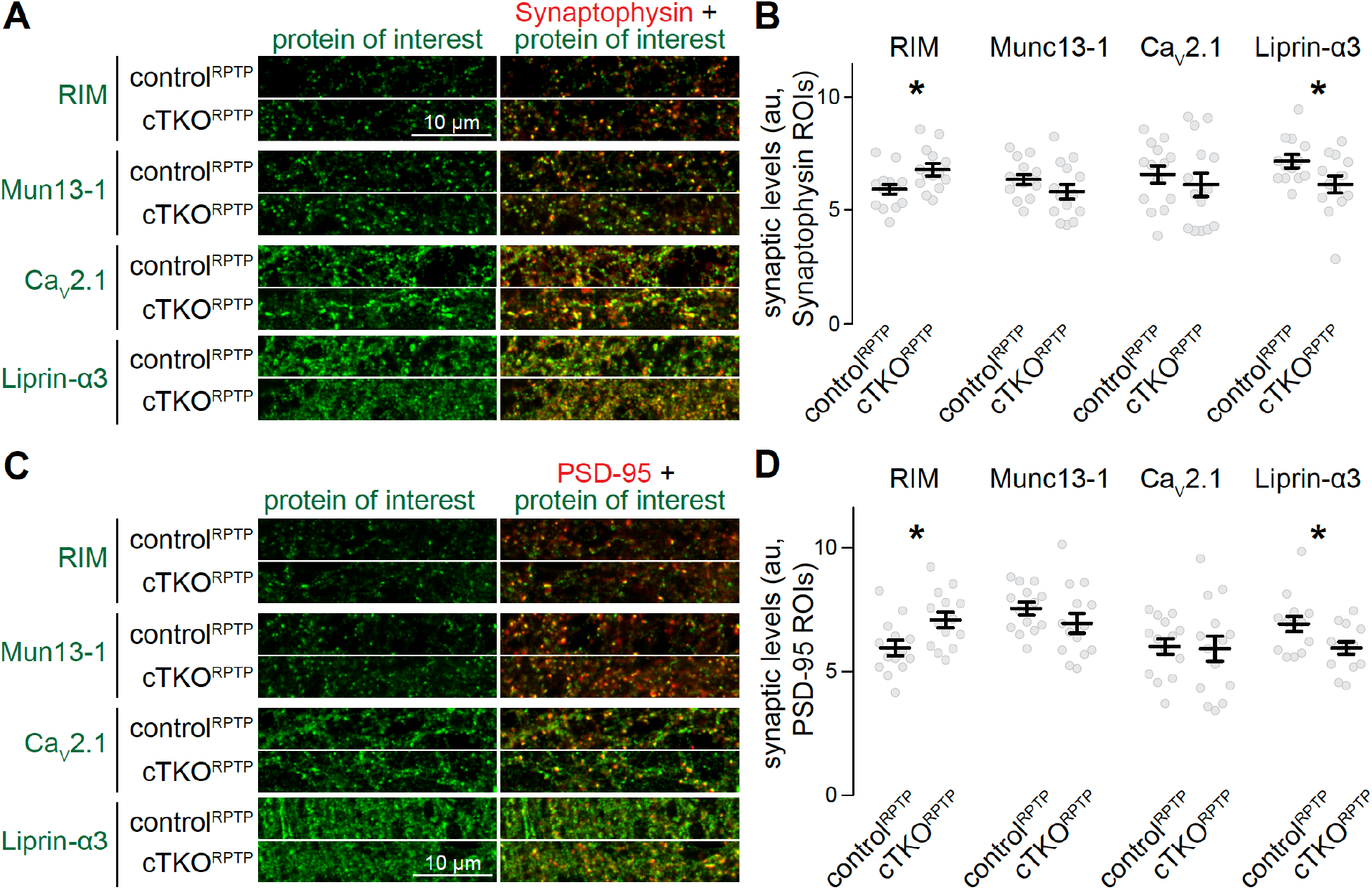
Confocal analyses of synaptic protein levels after ablation of LAR-RPTPs. **(A, B)** Example confocal images (A) and quantification (B) of the intensities of RIM, Munc13-1, Ca_V_2.1 and Liprin-α3 within Synaptophysin ROIs. The confocal images analyzed here were acquired in the same imaging session and for the same image frames as the STED analyses shown in Fig. 2. Confocal images were always acquired prior to STED acquisition, RIM: N (control^RPTP^) = 14 images/3 independent cultures, N (cTKO^RPTP^) = 14/3; Munc13-1: N (control^RPTP^) = 14/3, N (cTKO^RPTP^) = 14/3; Ca_V_2.1: N (control^RPTP^) = 14/3, N (cTKO^RPTP^) = 14/3; Liprin-α3: N (control^RPTP^) = 13/3, N (cTKO^RPTP^) = 13/3. **(C, D)** Same as A and B, but for PSD-95 ROIs. To avoid potential confounds of mildly increased Synaptophysin areas (Fig. 1M), we repeated the confocal analysis generating PSD-95 instead of Synaptophysin ROIs. In diffraction-limited microscopy, the resolution is insufficient to distinguish pre- and postsynaptic markers, and either marker can be used to generate synapse ROIs. N as in B. Data are plotted as mean ± SEM and were analyzed using t-tests, except for Ca_V_2.1 in B where a Mann-Whitney rank sum test was used. * p < 0.05.

## References

Ackley, B.D., Harrington, R.J., Hudson, M.L., Williams, L., Kenyon, C.J., Chisholm, A.D., and Jin, Y. (2005). The two isoforms of the Caenorhabditis elegans leukocyte-common antigen related receptor tyrosine phosphatase PTP-3 function independently in axon guidance and synapse formation. J. Neurosci. 25, 7517–7528.

Aicher, B., Lerch, M.M., Müller, T., Schilling, J., and Ullrich, A. (1997). Cellular redistribution of protein tyrosine phosphatases LAR and PTPsigma by inducible proteolytic processing. J. Cell Biol. 138, 681–696.

Biederer, T., Kaeser, P.S., and Blanpied, T.A. (2017). Transcellular Nanoalignment of Synaptic Function. Neuron 96, 680–696.

Bomkamp, C., Padmanabhan, N., Karimi, B., Ge, Y., Chao, J.T., Loewen, C.J.R., Siddiqui, T.J., and Craig, A.M. (2019). Mechanisms of PTPσ-Mediated Presynaptic Differentiation. Front. Synaptic Neurosci. 11, 17.

Bradley, A., Anastassiadis, K., Ayadi, A., Battey, J.F., Bell, C., Birling, M.-C., Bottomley, J., Brown, S.D., Bürger, A., Bult, C.J., et al. (2012). The mammalian gene function resource: the International Knockout Mouse Consortium. Mamm. Genome 23, 580–586.

Bunin, A., Sisirak, V., Ghosh, H.S., Grajkowska, L.T., Hou, Z.E., Miron, M., Yang, C., Ceribelli, M., Uetani, N., Chaperot, L., et al. (2015). Protein Tyrosine Phosphatase PTPRS Is an Inhibitory Receptor on Human and Murine Plasmacytoid Dendritic Cells. Immunity 43, 277–288.

Chen, L.Y., Jiang, M., Zhang, B., Gokce, O., and Sudhof, T.C. (2017). Conditional Deletion of All Neurexins Defines Diversity of Essential Synaptic Organizer Functions for Neurexins. Neuron 94, 611–625.e4.

Clandinin, T.R., Lee, C.H., Herman, T., Lee, R.C., Yang, A.Y., Ovasapyan, S., and Zipursky, S.L. (2001). Drosophila LAR regulates R1-R6 and R7 target specificity in the visual system. Neuron 32, 237–248.

Desai, C.J., Krueger, N.X., Saito, H., and Zinn, K. (1997). Competition and cooperation among receptor tyrosine phosphatases control motoneuron growth cone guidance in Drosophila. Development 124, 1941–1952.

Dunah, A.W., Hueske, E., Wyszynski, M., Hoogenraad, C.C., Jaworski, J., Pak, D.T., Simonetta, A., Liu, G., and Sheng, M. (2005). LAR receptor protein tyrosine phosphatases in the development and maintenance of excitatory synapses. Nat. Neurosci. 8, 458–467.

Dymecki, S.M. (1996). Flp recombinase promotes site-specific DNA recombination in embryonic stem cells and transgenic mice. Proc. Natl. Acad. Sci. U. S. A. 93, 6191–6196.

Elchebly, M., Wagner, J., Kennedy, T.E., Lanctôt, C., Michaliszyn, E., Itié, A., Drouin, J., and Tremblay, M.L. (1999). Neuroendocrine dysplasia in mice lacking protein tyrosine phosphatase σ. Nat. Genet. 21, 330–333.

Emperador-Melero, J., and Kaeser, P.S. (2020). Assembly of the presynaptic active zone. Curr. Opin. Neurobiol. 63, 95–103.

Emperador-Melero, J., Wong, M.Y., Wang, S.S.H., De Nola, G., Kirchhausen, T., and Kaeser, P.S. (2020). Phosphorylation triggers presynaptic phase separation of Liprin-α3 to control active zone structure. BioRxiv 2020.10.28.357574.

Farhy-Tselnicker, I., van Casteren, A.C.M., Lee, A., Chang, V.T., Aricescu, A.R., and Allen, N.J. (2017). Astrocyte-Secreted Glypican 4 Regulates Release of Neuronal Pentraxin 1 from Axons to Induce Functional Synapse Formation. Neuron 96, 428–445.e13.

Fukai, S., and Yoshida, T. (2020). Roles of type IIa receptor protein tyrosine phosphatases as synaptic organizers. FEBS J. febs.15666.

Han, K.A., Ko, J.S., Pramanik, G., Kim, J.Y., Tabuchi, K., Um, J.W., and Ko, J. (2018). PTPσ Drives Excitatory Presynaptic Assembly via Various Extracellular and Intracellular Mechanisms. J. Neurosci. 38, 6700–6721.

Han, K.A., Kim, Y.-J., Yoon, T.H., Kim, H., Bae, S., Um, J.W., Choi, S.-Y., and Ko, J. (2020a). LAR-RPTPs Directly Interact with Neurexins to Coordinate Bidirectional Assembly of Molecular Machineries. J. Neurosci. 40, 8438–8462.

Han, K.A., Lee, H.-Y., Lim, D., Shin, J., Yoon, T.H., Lee, C., Rhee, J.-S., Liu, X., Um, J.W., Choi, S.-Y., et al. (2020b). PTPσ Controls Presynaptic Organization of Neurotransmitter Release Machinery at Excitatory Synapses. IScience 23, 101203.

Held, R.G., Liu, C., Ma, K., Ramsey, A.M., Tarr, T.B., De Nola, G., Wang, S.S.H., Wang, J., van den Maagdenberg, A.M.J.M., Schneider, T., et al. (2020). Synapse and Active Zone Assembly in the Absence of Presynaptic Ca2+ Channels and Ca2+ Entry. Neuron 107, 667–683.e9.

Horn, K.E., Xu, B., Gobert, D., Hamam, B.N., Thompson, K.M., Wu, C.-L., Bouchard, J.-F., Uetani, N., Racine, R.J., Tremblay, M.L., et al. (2012). Receptor protein tyrosine phosphatase sigma regulates synapse structure, function and plasticity. J. Neurochem. 122, 147–161.

Johnson, K.G., and Van Vactor, D. (2003). Receptor Protein Tyrosine Phosphatases in Nervous System Development. Physiol. Rev. 83, 1–24.

Kaufmann, N., DeProto, J., Ranjan, R., Wan, H., and Van Vactor, D. (2002). Drosophila liprin-alpha and the receptor phosphatase Dlar control synapse morphogenesis. Neuron 34, 27–38.

Krueger, N.X., Van Vactor, D., Wan, H.I., Gelbart, W.M., Goodman, C.S., and Saito, H. (1996). The Transmembrane Tyrosine Phosphatase DLAR Controls Motor Axon Guidance in Drosophila. Cell 84, 611–622.

Kwon, S.-K., Woo, J., Kim, S.-Y., Kim, H., and Kim, E. (2010). Trans-synaptic Adhesions between Netrin-G Ligand-3 (NGL-3) and Receptor Tyrosine Phosphatases LAR, Protein-tyrosine Phosphatase δ (PTPδ), and PTPσ via Specific Domains Regulate Excitatory Synapse Formation. J. Biol. Chem. 285, 13966–13978.

Liu, C., Bickford, L.S., Held, R.G., Nyitrai, H., Sudhof, T.C., and Kaeser, P.S. (2014). The Active Zone Protein Family ELKS Supports Ca2+ Influx at Nerve Terminals of Inhibitory Hippocampal Neurons. J. Neurosci. 34, 12289–12303.

Liu, C., Kershberg, L., Wang, J., Schneeberger, S., and Kaeser, P.S. (2018). Dopamine Secretion Is Mediated by Sparse Active Zone-like Release Sites. Cell 172, 706–718.

McDonald, N.A., Fetter, R.D., and Shen, K. (2020). Assembly of synaptic active zones requires phase separation of scaffold molecules. Nature 588, 454–458.

Park, H., Choi, Y., Jung, H., Kim, S., Lee, S., Han, H., Kweon, H., Kang, S., Sim, W.S., Koopmans, F., et al. (2020). Splice-dependent trans-synaptic PTPδ-IL1RAPL1 interaction regulates synapse formation and non-REM sleep. EMBO J. 39, e104150.

Perez de Arce, K., Schrod, N., Metzbower, S.W.R., Allgeyer, E., Kong, G.K.-W., Tang, A.-H., Krupp, A.J., Stein, V., Liu, X., Bewersdorf, J., et al. (2015). Topographic Mapping of the Synaptic Cleft into Adhesive Nanodomains. Neuron 88, 1165–1172.

Prakash, S., McLendon, H.M., Dubreuil, C.I., Ghose, A., Hwa, J., Dennehy, K.A., Tomalty, K.M.H., Clark, K.L., Van Vactor, D., and Clandinin, T.R. (2009). Complex interactions amongst N-cadherin, DLAR, and Liprin-alpha regulate Drosophila photoreceptor axon targeting. Dev. Biol. 336, 10–19.

Pulido, R., Serra-Pages, C., Tang, M., and Streuli, M. (1995). The LAR/PTP delta/PTP sigma subfamily of transmembrane protein-tyrosine-phosphatases: multiple human LAR, PTP delta, and PTP sigma isoforms are expressed in a tissue-specific manner and associate with the LAR-interacting protein LIP.1. Proc. Natl. Acad. Sci. 92, 11686–11690.

Rizalar, F.S., Roosen, D.A., and Haucke, V. (2021). A Presynaptic Perspective on Transport and Assembly Mechanisms for Synapse Formation. Neuron 109, 27–41.

Sclip, A., and Südhof, T.C. (2020). LAR receptor phospho-tyrosine phosphatases regulate NMDA-receptor responses. Elife 9.

Serra-Pages, C., Medley, Q.G., Tang, M., Hart, A., and Streuli, M. (1998). Liprins, a family of LAR transmembrane protein-tyrosine phosphatase-interacting proteins. J. Biol. Chem. 273, 15611–15620.

Shishikura, M., Nakamura, F., Yamashita, N., Uetani, N., Iwakura, Y., and Goshima, Y. (2016). Expression of receptor protein tyrosine phosphatase δ, PTPδ, in mouse central nervous system. Brain Res. 1642, 244–254.

Skarnes, W.C., Rosen, B., West, A.P., Koutsourakis, M., Bushell, W., Iyer, V., Mujica, A.O., Thomas, M., Harrow, J., Cox, T., et al. (2011). A conditional knockout resource for the genome-wide study of mouse gene function. Nature 474, 337–342.

Südhof, T.C. (2012). The Presynaptic Active Zone. Neuron 75, 11–25.

Südhof, T.C. (2018). Towards an Understanding of Synapse Formation. Neuron 100, 276–293.

Takahashi, H., and Craig, A.M. (2013). Protein tyrosine phosphatases PTPδ, PTPσ, and LAR: presynaptic hubs for synapse organization. Trends Neurosci. 36, 522–534.

Takahashi, H., Arstikaitis, P., Prasad, T., Bartlett, T.E., Wang, Y.T., Murphy, T.H., and Craig, A.M. (2011). Postsynaptic TrkC and Presynaptic PTPσ Function as a Bidirectional Excitatory Synaptic Organizing Complex. Neuron 69, 287–303.

Uetani, N., Kato, K., Ogura, H., Mizuno, K., Kawano, K., Mikoshiba, K., Yakura, H., Asano, M., and Iwakura, Y. (2000). Impaired learning with enhanced hippocampal long-term potentiation in PTPdelta-deficient mice. EMBO J. 19, 2775–2785.

Um, J.W., and Ko, J. (2013). LAR-RPTPs: Synaptic adhesion molecules that shape synapse development. Trends Cell Biol. 23, 465–475.

Wallace, M.J., Batt, J., Fladd, C.A., Henderson, J.T., Skarnes, W., and Rotin, D. (1999). Neuronal defects and posterior pituitary hypoplasia in mice lacking the receptor tyrosine phosphatase PTPsigma. Nat. Genet. 21, 334–338.

Wang, S.S.H., Held, R.G., Wong, M.Y., Liu, C., Karakhanyan, A., and Kaeser, P.S. (2016). Fusion Competent Synaptic Vesicles Persist upon Active Zone Disruption and Loss of Vesicle Docking. Neuron 91, 777–791.

Wong, M.Y., Liu, C., Wang, S.S.H., Roquas, A.C.F., Fowler, S.C., and Kaeser, P.S. (2018). Liprin-α3 controls vesicle docking and exocytosis at the active zone of hippocampal synapses. Proc. Natl. Acad. Sci. 115, 2234–2239.

Wu, X., Cai, Q., Shen, Z., Chen, X., Zeng, M., Du, S., and Zhang, M. (2019). RIM and RIM-BP Form Presynaptic Active-Zone-like Condensates via Phase Separation. Mol. Cell 73, 971–984.e5.

Yang, T., Bernabeu, R., Xie, Y., Zhang, J.S., Massa, S.M., Rempel, H.C., and Longo, F.M. (2003). Leukocyte Antigen-Related Protein Tyrosine Phosphatase Receptor: A Small Ectodomain Isoform Functions as a Homophilic Ligand and Promotes Neurite Outgrowth. J. Neurosci. 23, 3353–3363.

Yim, Y.S., Kwon, Y., Nam, J., Yoon, H.I., Lee, K., Kim, D.G., Kim, E., Kim, C.H., and Ko, J. (2013). Slitrks control excitatory and inhibitory synapse formation with LAR receptor protein tyrosine phosphatases. Proc. Natl. Acad. Sci. 110, 4057–4062.

Zucker, R.S., and Regehr, W.G. (2002). Short-term synaptic plasticity. Annu. Rev. Physiol. 64, 355–405.

